# Geometrical Designs in Volumetric Bioprinting to Study Cellular Behaviors in Engineered Constructs

**DOI:** 10.1101/2025.07.14.664683

**Authors:** Julia Simińska-Stanny, Pierre Tournier, Armin Shavandi, Shukry J Habib

**Affiliations:** Université libre de Bruxelles (ULB), École Polytechnique de Bruxelles, 3BIO-BioMatter, Avenue F.D. Roosevelt, 50 - CP 165/61, 1050 Brussels, Belgium; Department of Biomedical Sciences, University of Lausanne, Switzerland

**Keywords:** volumetric printing, curvature, perfusion, vascular models, composite materials

## Abstract

This study investigates the influence of geometrical variations in volumetrically printed (Vol3DP) structures on the attachment, survival, and organization of cancer cells (143b) and human umbilical vein endothelial cells (HUVECs). We adapted a gelatin methacryloyl (GelMA)–poly(ethylene glycol) diacrylate (Gel-PEG) resin for volumetric bioprinting. Compared to GelMA, Gel-PEG improved printing fidelity and resolution, superior mechanical properties, and reduced swelling. We fabricated disc-like constructs and channel geometries, including straight channels and angular designs of 60°, 90°, and 110° and cultured human umbilical vein endothelial cells (HUVECs) and 143b human osteosarcoma cells, a highly metastatic cell line, for up to 14 days. Using label-free holographic microscopy, we visualized cellular protrusions, important for adhesion and mechanosensing, in real-time and without staining, an advantage for long-term, live-cell analysis in 3D constructs. HUVECs adhered well, expressed CD31, and showed preferential spreading in channels with specific geometrical angles, indicating geometry-sensitive behavior. This is physiologically relevant, as it reflects the native mechanosensitive and alignment behavior of endothelial cells during vascular formation. In contrast, osteosarcoma cells spread uniformly throughout the constructs, formed dense, geometry-independent agglomerates, and exhibited enhanced growth and spreading within the Gel-PEG matrix compared to GelMA. This behavior is consistent with the aggressive and geometry-insensitive nature of metastatic tumor cells. These findings highlight Gel-PEG’s utility for generating stable, biomimetic 3D environments and demonstrate the application of holographic microscopy for assessing cell– material interactions within volumetrically bioprinted constructs, underscoring the potential of this approach for developing vascularized models and studying mechanobiological responses in engineered tissues.

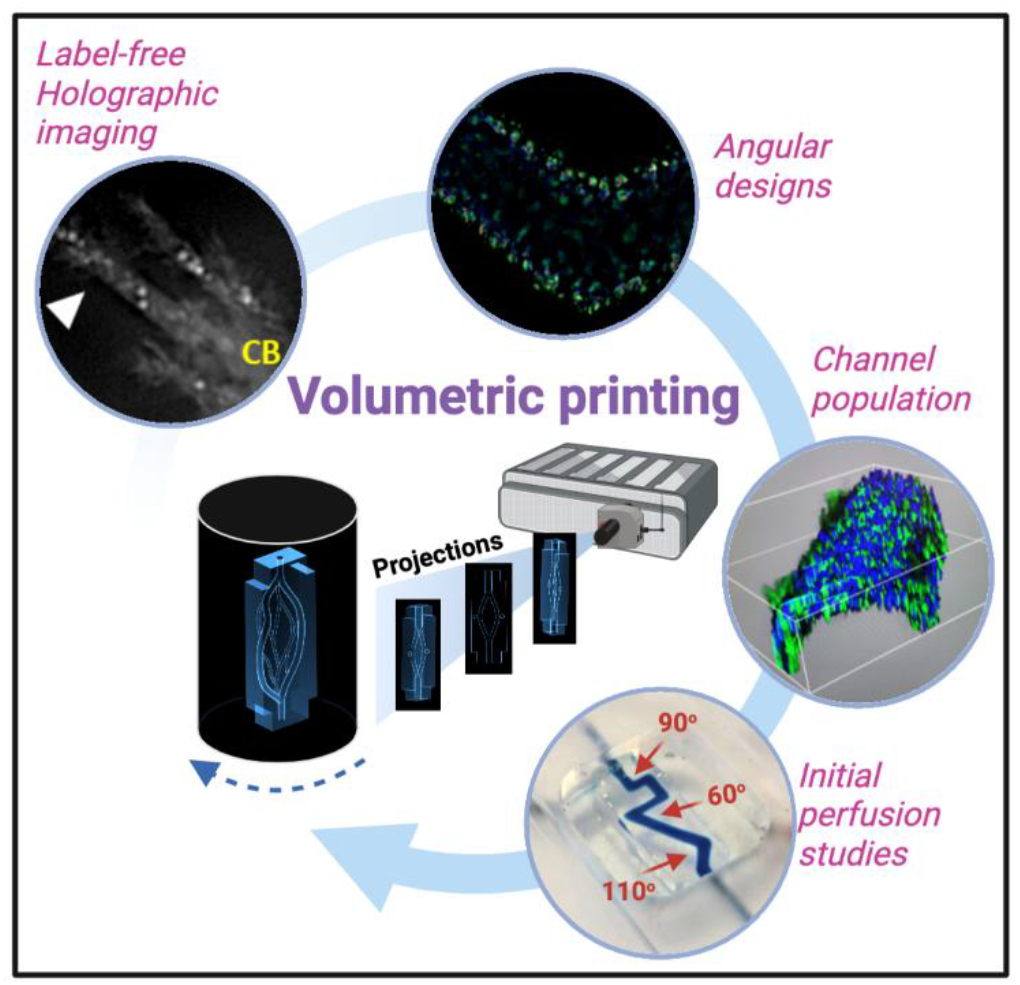

## 1. Introduction

The engineering of three-dimensional (3D) tissue models that recapitulate in vivo cellular microenvironments is a critical goal in regenerative medicine, drug screening, and fundamental cell biology [1]. Traditional two-dimensional (2D) cell culture systems fail to mimic the complex architecture, biochemical cues, and mechanical properties present in native tissues, limiting their physiological relevance and predictive value [2]. Volumetric bioprinting has recently emerged as a transformative technique, enabling rapid fabrication of intricate 3D constructs with high spatial resolution and scalability, thus offering opportunities for creating biomimetic tissue models [3-5]. Among key biophysical cues regulating cell behavior, substrate curvature plays a central role in modulating adhesion, migration, proliferation, and differentiation [6]. High-curvature regions typically enhance cell proliferation and motility, while low-curvature areas promote alignment and elongation, particularly in mechanosensitive endothelial cells [3-5]. These responses involve cytoskeletal tension and nuclear deformation, with the nucleus sensing curvature and translating mechanical forces into changes in gene expression [1, 7]. Such mechanotransduction processes are crucial for guiding cell organization and fate, impacting scaffold design in tissue engineering. Callens and coworkers reported that on convex surfaces, fibroblasts and mesenchymal stem cells align longitudinally to minimize bending energy, while epithelial cells align circumferentially due to stress fiber contractility. Spherical surfaces inhibit cell motility, while concave microstructures promote rapid migration. Saddle surfaces, with both convex and concave curvatures, induce cell alignment in orthogonal directions [1]. Simulations of cell adhesion and migration on sinusoidal surfaces demonstrate that 3D curvature induces intracellular pressure gradients [2]. Curvature can act as a potent regulator of cell fate. High-curvature regions often promote cell proliferation and migration, while low-curvature areas may enhance alignment and elongation, particularly in endothelial cells [8]. This interaction between curvature and cellular behavior has significant implications for tissue engineering, where precise control over cell fate and tissue architecture is crucial for functional tissue regeneration.

While “vessel-on-chip” systems have emerged as valuable tools for studying endothelial cells in physiologically relevant environments, the impact of substrate curvature, a critical feature of the *in vivo* vasculature, has been largely underscored in these models [9]. The ability to replicate physiological microenvironments through 3D bioprinting has transformed the field of tissue engineering. Among the diverse bioprinting techniques, volumetric bioprinting has emerged as a method of rapidly fabricating intricate biomimetic structures with high spatial resolution [10-12]. Understanding how cells respond to specific geometrical and material cues within their microenvironment remains a major challenge in tissue engineering and regenerative medicine [15]. Endothelial cells, such as human umbilical vein endothelial cells (HUVECs), are mechanosensitive and highly responsive to geometric cues, which regulate their adhesion, spreading, and organization into vascular networks [6, 16, 17]. Similarly, tumor cells like the highly metastatic 143b osteosarcoma line exhibit aggressive and geometry-independent growth patterns that can be exploited to model cancer progression in engineered microenvironments [18, 19]. Advanced imaging techniques that allow real-time, label-free monitoring of cell-material interactions in 3D constructs are crucial for elucidating these behaviors [15]. Holographic microscopy, offering quantitative phase imaging without the need for exogenous labels, has proven valuable in 2D and simple 3D culture systems [20]. However, its application to complex volumetrically bioprinted constructs is still in its infancy. Despite significant advancements in 3D bioprinting and tissue engineering, there remains a need for a deeper understanding of how specific geometrical features influence cell-material interactions and ultimately dictate tissue-level outcomes. Studying these dynamics in well-defined hierarchical 3D structures is essential for designing biomaterials that can support functional tissue regeneration or provide accurate disease models. This study aims to provide insights into the spatial organization and functional behavior of human cancer cells (143b osteosarcoma cell line) and human umbilical vein endothelial cells (HUVECs) within volumetrically printed bioresin constructs that vary in printability, mechanical properties, stability profiles, and geometrical features.

## 2. Results and discussion

### 2.1 Enhanced Printability and Mechanical Stability of GelMA–PEGDA Composite Resins for Volumetric Bioprinting

Composite materials are surpassing conventional ones providing better tunability and easier processing [10, 21], but their use in volumetric printing remains underexplored. Here we combined methacryloyl groups grafted to gelatin (GelMA) with synthetic polyethylene glycol diacrylate (PEGDA) polymer for simultaneous light-triggered crosslinking and the creation of hybrid network composite. The composite Gel-PEG resin showed suitability for volumetric printing (**Figure 1A**) due to enhanced transparency (**Figure 1B**), allowing for the precise generation of the object geometry within the photoresin. Gel-PEG resin eased printed object detection (**Figure 1C-ii**) compared to GelMA resin (**Figure 1C-i**), once the process is completed. Enhanced polymerization control allowed by Gel-PEG translated for precisely cured unobstructed channel geometries capable of perfusion (**Figure 1C, Supplementary Information (SI) – video S2, S3**).

**Figure 1.**
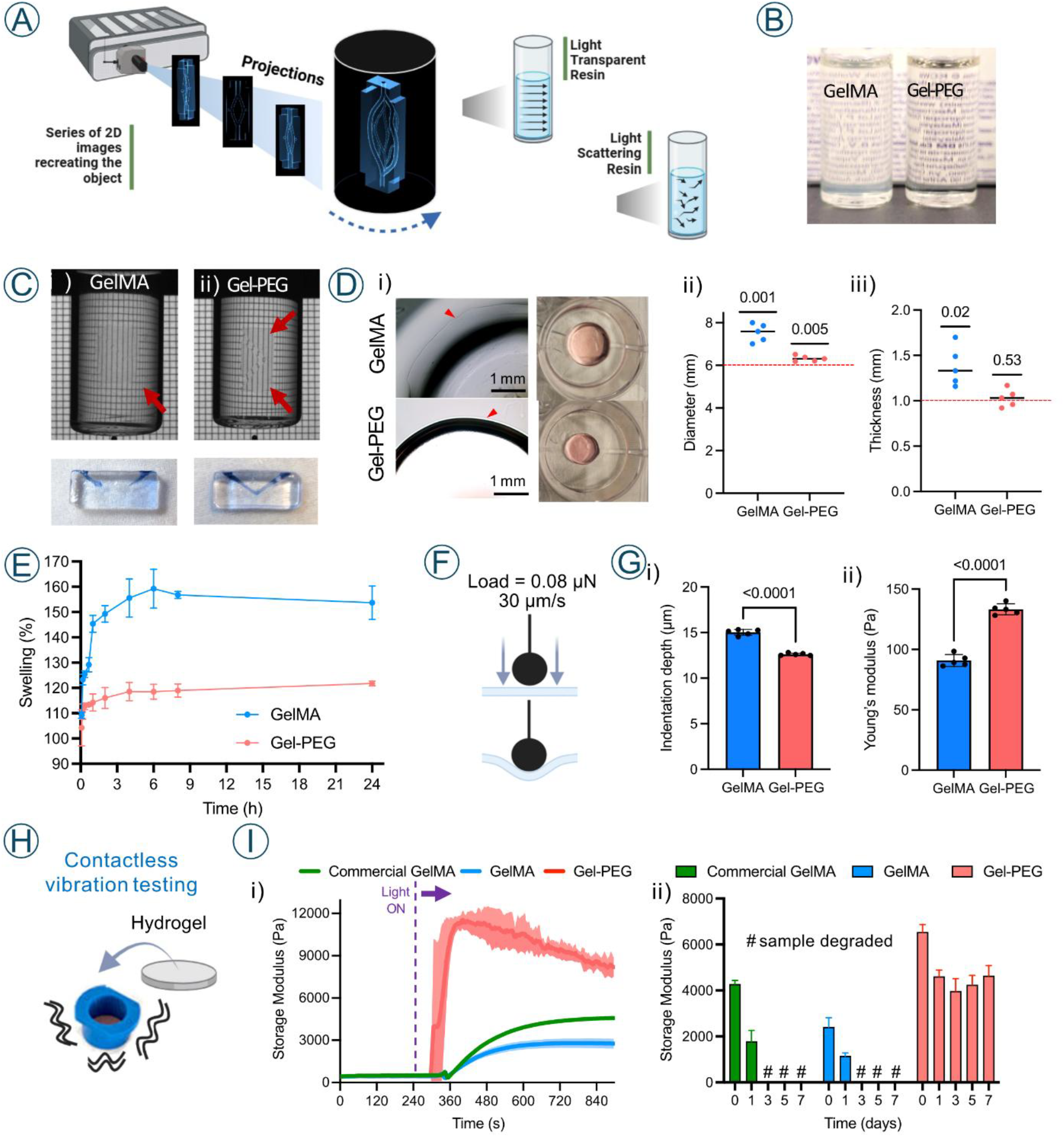
**A)** Schematic of volumetric printing of double-spiral channeled geometry, with the comparison of how the light travels in transparent and scattering resins; **B)** GelMA (left) and Gel-PEG (right) resins loaded into printing vials; **C)** Comparison of the volumetric printing results for GelMA (i) and Gel-PEG (ii) resins, the printed shape can be easily distinguished for Gel-PEG resin and channel patency assessment for the constructs fabricated with GelMA (i) and Gel-PEG (ii) performed by perfusing blue-colored PBS through the printed channel; **D) (i)** Representative brightfield images of GelMA and Gel-PEG hydrogels after volumetric 3D printing (left), red arrowheads indicate the edges of the hydrogels, and after 24 h incubation in cell medium (right), (ii-iii) Quantification of hydrogel dimensions showings that Gel-PEG samples more closely match the theoretical dimensions (i.e., digital input, red dashed line: 6 mm diameter × 1 mm thickness) compared to GelMA. n = 5 gels per group. Statistical test: one sample t-test against the theoretical mean. **E)** Swelling test for GelMA and Gel-PEG hydrogels, based on the mass differences at different time points n=3 gels per group. **F)** Schematic of the mechanical indentation setup used to measure stiffness, applying a 0.08 μN load at a rate of 30 μm/s **G)** (i) Indentation testing of hydrogels: Gel-PEG hydrogels showed significantly reduced indentation depth compared to GelMA, (ii) Young’s modulus (stiffness) of 3D-printed GelMA and Gel-PEG hydrogels. N = 5 gels per group, n ≥ 16 measurements per gel. Statistical test: Welch’s t-test. **H)** Schematics of the nondestructive vibrational hydrogel testing, where the hydrogel sample is formed directly on the membrane within a holder (blue). **I)** (i) Photopolymerization tests for Gel-PEG, home-made GelMA and commercial GelMA, showing the changes of storage modulus in time (ii) and the values of storage modulus of Gel-PEG, GelMA and commercial GelMA measured at different time points following samples incubation in PBS. All data are presented as mean ± standard deviation.

Gel-PEG hydrogels exhibited improved, defined edges (**Figure 1D-i**) and 3D printing fidelity, with diameters and thicknesses more closely matching the Computer Aided Design (CAD) model than those of GelMA samples (**Figure 1D-ii,iii, Figure S1.1**). Gel-PEG hydrogels also demonstrated superior structural stability with minimal swelling over time compared to GelMA constructs, which developed uneven surfaces. Quantitative swelling analysis revealed that GelMA samples absorbed more than twice the amount of water relative to Gel-PEG discs (**Figure 1E**).

Indentation testing indicated significantly lower indentation depths for Gel-PEG hydrogels (**Figure 1G-i**), despite similar applied peak loads across both groups (**Figure S1.1**). Young’s modulus measurements confirmed enhanced mechanical strength of Gel-PEG constructs (∼130 Pa) compared to GelMA (∼90 Pa) (**Figure 1F, 1G-ii**).

Complementary vibrational viscoelasticity tests (**Figure 1H**) supported these findings, showing distinct mechanical profiles and polymerization kinetics, assessed via time-dependent changes in storage modulus (G’). The results revealed that Gel-PEG hydrogels rapidly reached a peak G’ of 12 kPa, which stabilized at ∼9 kPa within 1–2 minutes (**Figure 1I-i**). In contrast, GelMA hydrogels reached a lower peak of G’ of 3 kPa, below that of the commercial GelMA benchmark, likely due to a reduced degree of methacrylation in the in-house formulation [22]. Time-course monitoring of viscoelastic properties over one week showed that GelMA samples experienced a two-fold decrease in G’ after one day and fully disintegrated by day three. In contrast, Gel-PEG hydrogels showed a modest reduction in G’ from ∼6.5 kPa to ∼4 kPa, which then remained stable (**Figure 1I-ii**), indicating sustained integrity and mechanical properties [23]. The improved performance of Gel-PEG aligns with studies indicating that the inclusion of PEGDA in the resin formulation offers superior mechanical properties and stability [24]. Additionally, this composition’s low swelling properties mitigate uncontrolled degradation, a common limitation in single-polymer systems [25, 26]. Such enhanced stability allows for consistent cell adhesion, crucial for long-term experiments.

### 2.2 Evaluating Cell Adhesion and Spreading on Volumetrically Printed GelMA–PEGDA constructs

To test the ability of the Gel-PEG constructs to support cell attachment and growth, we seeded primary human umbilical vascular endothelial cells (HUVECs) or human cancer cells from the 143b osteosarcoma cell line on the printed disks (**Figure 2A**), and we first evaluated their ability to adhere to the surfaces.

**Figure 2.**
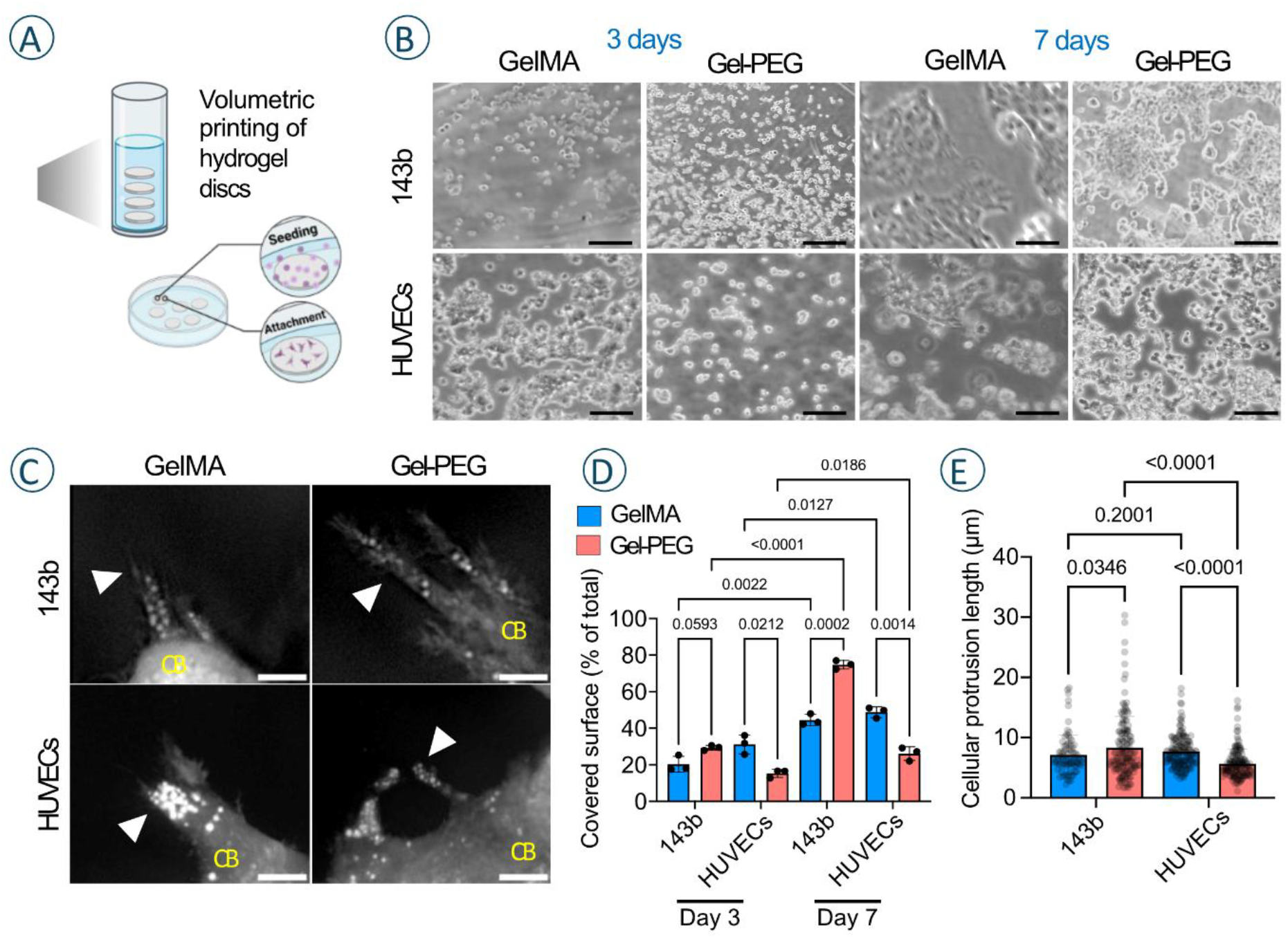
**A)** Schematic of the cell seeding on the discs to evaluate cell attachment and growth; **B)** Representative images of HUVECs seeded on the discs after 3 and 7 days in culture. The scale bar corresponds to 0.5 mm; **C)** Representative holographic microscopy images of **(C)** CB: cell body, white arrowhead: protrusions, scale bars: 0.5 µm. All data are presented as mean ± standard deviation. Statistics: Unpaired t with Welch’s correction; **D)** Quantification of the area covered by cells after 3 and 7 days of culture for 143b cells and HUVECs on the disks. n = 3 random areas per disk; **E)** Quantification of cellular protrusion length for 143b cells and HUVECs on the disks, (n ≥ 83 protrusions per group, taken from ≥ 10 images/group).

As seen in **Figure 2B**, both cell types were able to adhere on Gel-PEG or GelMA surfaces, likely due to the presence of gelatin, rich in cell adhesion motifs [27], highlighting the suitability for Gel-PEG bioresin for further use in tissue engineering. After 3 days, the colonization of GelMA or Gel-PEG disks was similar for the osteosarcoma cells, whilst HUVECs preferentially adhere on GelMA. After 7 days, both hydrogels supported cell growth, yet with material-dependent cell proliferation with the same observation at day 3 (**Figure 2C**). As cellular adhesion and motility require cytoskeletal reorganization and engagement of cellular protrusions, we used label-free holographic imaging of live cells (**Figure 2D, Figure S1.2**) to reveal the spreading of cells and the formation of cellular extensions and adhesion points in both GelMA and Gel-PEG surfaces. This observation showed material-and cell-dependent response, as cellular protrusions were longer on Gel-PEG compared to GelMA for 143b cells, while the opposite effect was observed for HUVECs (**Figure 2E**). Interestingly, we also observed a mobilization of lipid droplets located in the distal parts of the cellular extensions (**Figure 2E**). By reorganizing their local cytoskeleton network, cells provide energy in specific subcellular regions with high demands. Thus, local lipid accumulation can modulate cell adhesion and invasion of the substrate material [28, 29].

These observations highlight the substrate-specific cell adhesion and underscore the need to design advanced specialized materials for tissue engineering applications.

### 2.3 Impact of Channel Geometry on Cellular Organization

Volumetric printing allows fast (<60 s) and easy shaping of hydrogel substrates into channeled designs [21], especially compared to cumbersome methods such as patterning microtunnels inside gelatin-gel blocks by photo-thermal etching [30] or multiple-step fabrication of microfluidics platforms [31]. The use of thermo- and photo-crosslinkable materials allows for recovering the printed object once the thermoresponsive matrix liquifies and also allows the reuse of the non-crosslinked material for successive prints (**Figure 3A**) [21, 32]. The composite resin used in this study had a similar photopolymerization pattern to widely used GelMA (**Figure 3B**) and allowed fabricating high model fidelity structures.

**Figure 3.**
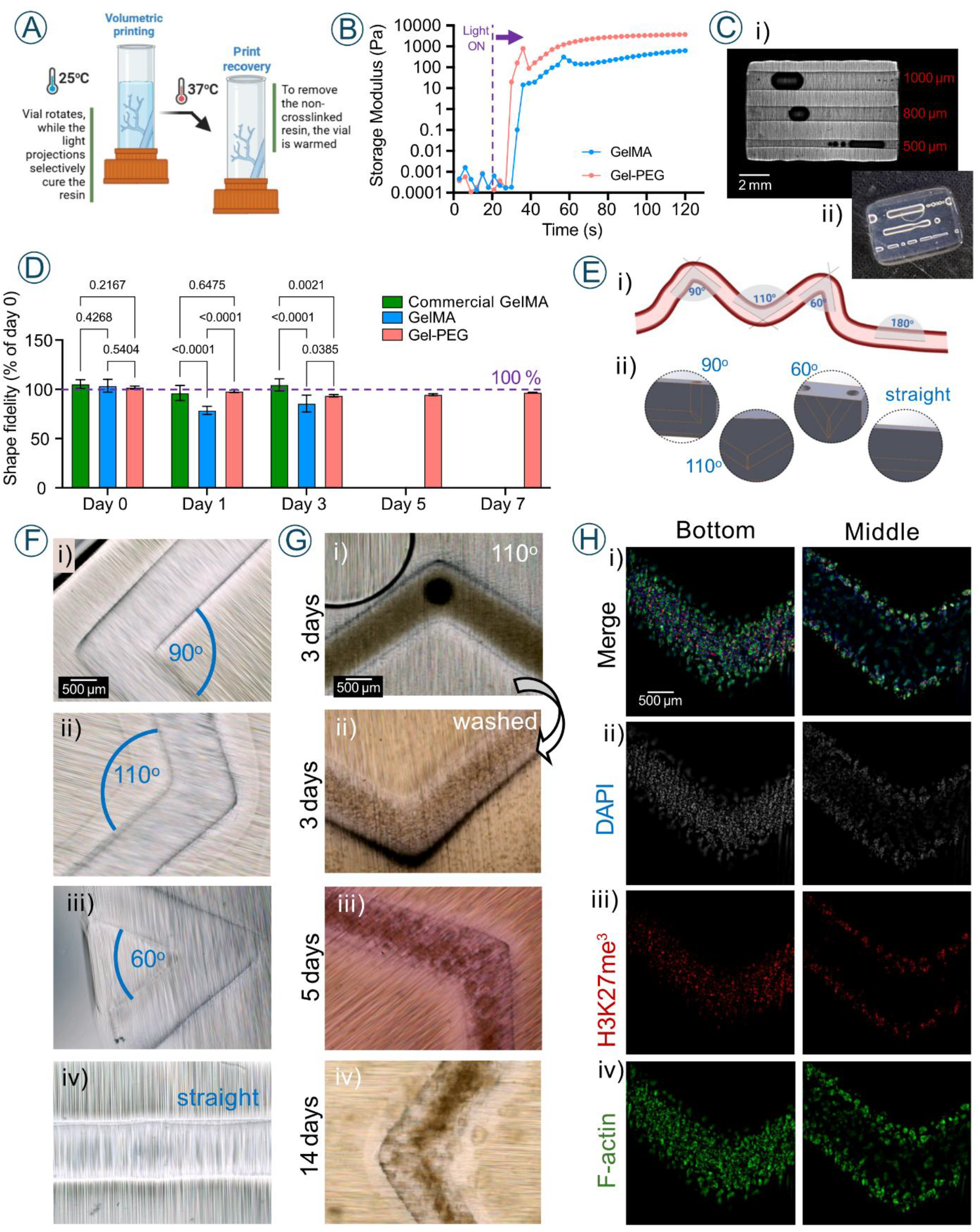
**A)** Principle of volumetric printing with thermogelling resins. Printing is performed at room temperature when the material is in gel state. After printing, the resin is warmed to release the construct and reuse non-crosslinked resin. **B)** Time-sweep test showing the photocrosslinking pattern for GelMA and Gel-PEG resins. The dashed line indicates the moment from which the 405 nm light was on. Representative plots, n=2. **C)** Volumetrically printed 3D model with varying diameter channels (500-1000 µm) to benchmark the material printability and shape fidelity (i) Coherence contrast microscopy image, (ii) photograph, **D)** Shape fidelity between the as-printed channel diameter and the diameter of the channels up to 7 days of incubation in PBS at 37 °C. n≥3, mean ± SD., one-way ANOVA with Dunn-Sidak post-hoc test. **E)** Schematic representation of vascular structure and its curvature (i) and (ii) corresponding CAD designs used to recapitulate native vessels curvature by straight and angled designs (60°, 90°, 110°). **F)** Volumetrically printed angled 90° (i), 110° (ii), 60° (iii) and straight (iv) geometrical designs. **G)** 143b cells seeded in 110° volumetrically printed channels; (i) cells proliferate within the channel, (ii) after 3 days the non-attached cells are washed, (iii) after 5 days cells form clusters and the following days in culture led to the increased number of cells at the bottom (iv - day 14). **H)** Immunofluorescence staining images (day 7) showing 143b cells that colonized the volumetrically printed channel. (i) Merged channels and individual channels of (ii) cell nuclei (DAPI, grey), (iii) Lysine 27 tri-methyl Histone H3 (red), and (iv) F-actin (green).

To evaluate the shape fidelity, we designed a multi-channel construct with three parallel channels of 500 µm, 700 µm, and 1000 µm (**Figure 3Ci-ii**). Constructs printed with the Gel-PEG resin exhibited high fidelity and stability, with only a ∼5% decrease in channel diameter observed after day 3, which remained stable through day 7, indicating good long-term structural stability in physiological conditions of osmolarity and temperature (i.e., PBS at 37 ^°^C). In contrast, GelMA-based constructs, both the commercial variant (60% degree of substitution) and in-house synthesized version (40%) showed varied performance. The commercial GelMA maintained good precision, with diameter changes remaining below 5% throughout the testing period. However, the 40% substituted GelMA displayed significant swelling, with up to 20% variation in channel size between day 0 and days 1–3. This swelling likely led to expansion of the hydrogel matrix and partial occlusion of the channels. Unlike the Gel-PEG resin, the 40% GelMA was unable to support printing of the smallest channel, and the middle channel was often non-perfusable. Furthermore, both GelMA variants lost structural integrity after 3 days, underscoring their limited stability compared to Gel-PEG (**Figure 3C-iii**). It is important to note that degradation observed in the in-house GelMA may be formulation-dependent, and results may not extrapolate to all GelMA hydrogels.

Collective cell migration, a crucial process for various biological functions such as cancer progression, metastasis, angiogenesis, or wound healing, involves the coordinated movement of cell cohorts. Recent research highlights how the three-dimensional topographical confinements significantly influence how cell groups move within confined spaces like tubular geometries [30]. The impact of substrate curvature on cells has only recently been investigated and is now recognized as a critical cue regulating cell behavior. In the vasculature, endothelial cells encounter curvature at the subcellular, cellular, and tissue scales [7, 9]. In this study, we chose to investigate angular designs of 60°, 90°, and 110° as well as straight grooves and accordingly designed simplified CAD models (**Figure 3D**). Various printed geometries (**Figure 3F**) were first seeded with 143b cells to optimize the cell seeding protocol. The highest seeding success was observed when cells were injected into the channels, incubated, and then, after 6-8 h, the constructs were flipped to facilitate cell attachment on the opposite side. After 3 days, the channels were washed with PBS (pre-warmed at 37^°^C) to remove the non-adherent cells (**Figure 3G-i, ii**). At longer time points (5 and 14 days), osteosarcoma 143b cells created agglomerates and densely populate the bottom of the channel (**Figure 3G-iii, iv**) causing slight stenosis of the channel. A similar phenomenon was previously described for cancer cells adhering to the endothelial layer lining a perfused channel [4].

After 7 days of culture, cancer cells achieved complete coverage of the channels (**Figure 3H**). DAPI-stained nuclei revealed significant cellular clustering, demonstrating their strong adaptability to the substrate’s topography. The observed nuclear clustering patterns were consistent with high proliferation rates typically seen in dense tumor models [33]. Staining of nuclei (DAPI (gray)) and cytoskeleton (Actin (green)) after 7 days in various angular confinements showed dense cell populations across all angles, underscoring the invasive and highly adaptable behavior of cancer cells. Immunofluorescence analysis of lysine 27 trimethylation of core histone 3 (H3K27me^3^), a histone modification associated with gene repression and implicated in cancer progression and therapy response, highlighted functional properties of 143b osteosarcoma cells within the constructs (**Figure 3H-iii, SI - Figure S1.3**).

We then sought to investigate how HUVECs could colonize vascular-like construct with various angles generated using volumetric 3D printing. After the first 3 days of culture, we observed that most HUVECs jammed on the bottom of the channels, but longer culture (5 days) demonstrated that HUVECs preferentially migrated to the channel edges and formed clusters (**Figure 4A**). HUVECs displayed a typical rounded morphology, and did not exhibit aberrant overgrowth as cancer cells, reminiscent to that of healthy (non-cancerous) cells. Immunofluorescence staining for platelet endothelial cell adhesion molecule (PECAM-1/CD31), an expression marker of human endothelial cells involved in adhesion, angiogenesis, and vascular integrity, confirmed the presence of HUVECs within the vascular-like constructs (**Figure 4A-C**).

**Figure 4.**
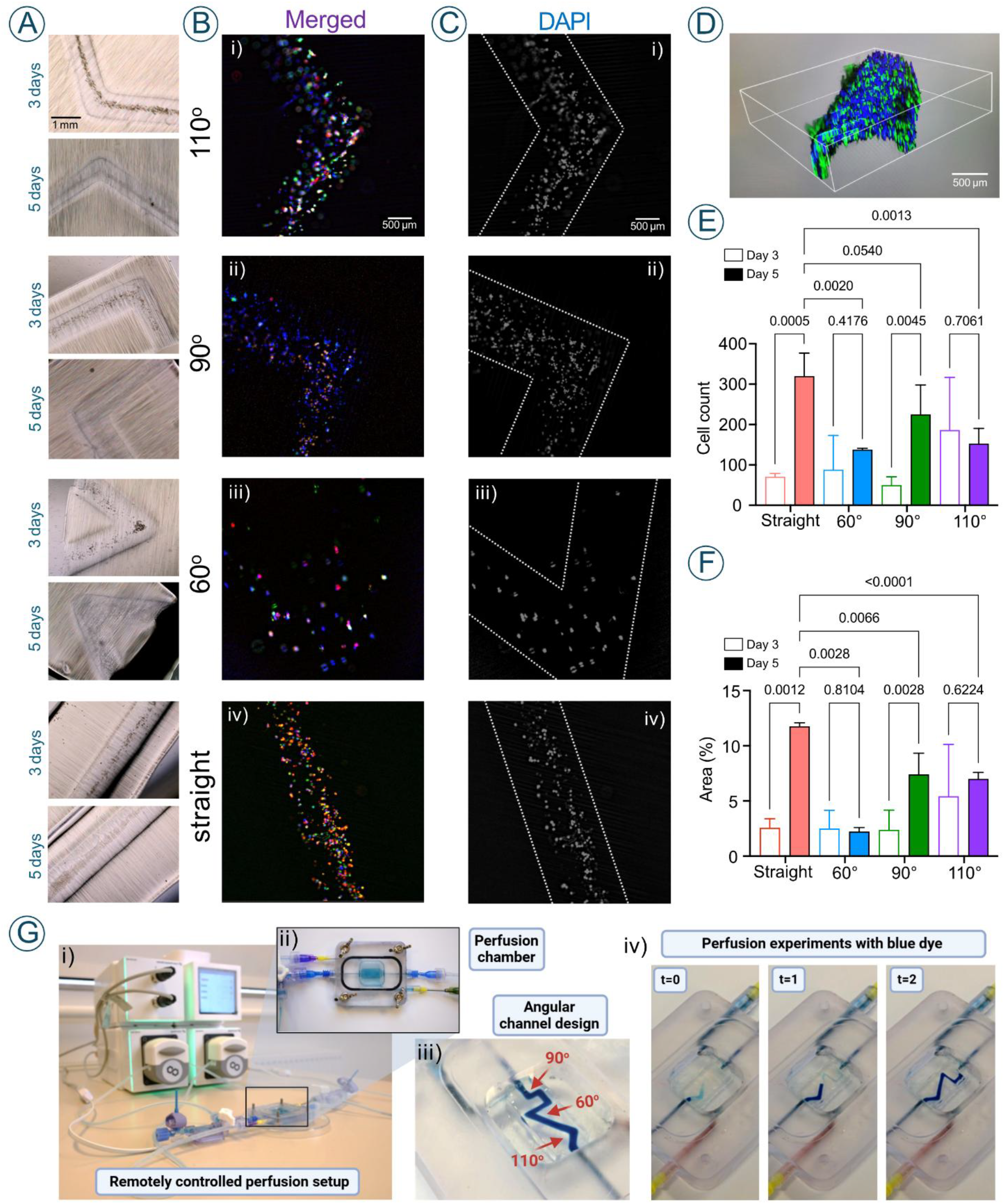
**A)** Representatives brightfield images of HUVECs seeded within different geometrical designs after 3 and 5 days in culture. **B-C**) Representatives, images of immunostaining of HUVECs seeded in volumetrically printed channels after 5 days in culture: **B)** Representative merged images showing the cell nuclei stained in grey (DAPI), cytoskeleton (F-actin) stained in green (Phalloidin-AF488) and PCAM-1/CD31 in red.; **C)** Representative merged images showing the cell nuclei stained in grey (DAPI), to assess cell distribution. **D)** Representative 3D reconstructed image of channel curvature based on z-stacked images. **E)** Number of cells counted in the different curvatures after 3 and 5 days of culture (n≥3, except day 5 60°, where n=2), mean ± SD, two-way ANOVA with Dunn-Sidak post-hoc test. **F)** Area covered by cell nuclei within the channeled geometries in different curvatures after 3 and 5 days of culture (n≥3, except for day 5, 60°, where n=2), mean ± SD, two-way ANOVA with Dunn-Sidak post-hoc test. **G)** Remotely controlled perfusion system (i) featuring a DLP-fabricated perfusion chamber (ii) designed for the dynamic evaluation of 3D-printed constructs.

Geometrical variations in the printed channels (straight, 60°, 90°, and 110°) had a pronounced effect on the HUVECs’ behavior. Cell density and occupied area were higher in 110° channels compared to 90°, 60° and straight designs after 3 days of culture (**Figure 4D-F**). Further quantitative analysis indicated that channels with acute (60°) or straight (90°) angles as well as longitudinal geometries resulted in an increased number of HUVECs in a 5-day culture compared to 3 days (**Figure 4E**). Conversely, acute angles (60°) resulted in uneven cell distributions and limited expansion understood as the area coverage of the channels, compared to straight (90°) angles as well as longitudinal geometries (**Figure 4F**).

Finally, as a proof of concept, we volumetrically printed a hydrogel with an angular channel and perfused it with methylene blue, demonstrating the system’s potential for simultaneous testing of various angular designs under dynamic flow conditions (**Figure 4G, video S4**). This capability is beneficial for modeling vascular flow dynamics, allowing investigation of how different channel angles influence fluid perfusion, shear stress, and nutrient delivery within engineered vascular networks. It also enables optimization of scaffold geometries to promote optimal flow and cell behavior, critical for tissue engineering and organ-on-chip applications. Furthermore, dynamic perfusion through complex geometries facilitates drug screening under physiologically relevant flow conditions and supports mechanobiological studies by revealing how combined fluid shear and substrate curvature impact cell adhesion, migration, and mechanosensing in 3D constructs.

It has been reported that acute angles can lower the persistence of cell migration [3]. For instance, work by Takai et. al., on angular biodevices demonstrated that different cell migration behavior stems from directional extensions and adhesion phenomena of each cell type [3]. Our findings corroborate these results, showing that acute (60°) channels caused a decrease in the number and area coverage of HUVECs, conversely to obtuse angles and straight geometries. However, these findings can likely be influenced by introducing flow conditions as previously evidenced to be a crucial factor in endothelial cells response [4, 5].

Overall, we observed that 143b cancer cells and HUVECs exhibited distinct behaviors within the same topographical confinements. Cancer cells demonstrated higher adaptability, forming dense agglomerates across channeled geometries. These differences align with studies emphasizing the invasive nature of cancer cells. Research by de Visser and Joyce highlighted that cancer cells can adapt to varied microenvironments by altering their adhesion properties [33]. Conversely, endothelial cells rely on stable junctions and nutrient availability, making them more sensitive to geometric and material constraints.

## 3. Conclusions

This study demonstrates the utility of volumetric bioprinting in creating complex microenvironments for studying cellular responses to geometric and mechanical cues, and substrate material composition. Our findings highlight the critical influence of geometrical features on cell adhesion, morphology, and functional behavior. Straight channels or with moderate angulation (90°-110°) were particularly effective in promoting HUVECs clustering and proliferation, while cancer cells displayed high adaptability across all geometries. The Gel-PEG resin proved to exhibit superior structural integrity, enhanced mechanical properties, and long-term stability, suitable for cell culture, compared to conventional materials, supporting robust cell attachment and long-term viability. Distinct differences between cancer cells and endothelial cells within these engineered constructs underscore the importance of tailored biomaterial and geometric design in tissue engineering and disease modeling.

The ability to fabricate and perfuse angular microchannels opens new avenues for modeling physiologically relevant vascular environments, facilitating investigations into how substrate curvature and flow dynamics influence cell behavior. In future studies, integrating a bioreactor can be used to enhance nutrient and oxygen delivery within constructs. It is also beneficial to evaluate the effects of sustained culture on cellular organization and functionality for a longer period. Finally, mechanobiological studies can be considered to investigate the impact of mechanical forces and substrate stiffness on cellular behavior.

## 4. Experimental section

### 4.1 Materials

Gelatin from porcine skin, phosphate-buffered saline (PBS), Methacrylic anhydride (MAA), polyethylene glycol diacrylate (PEGDA, average Mn 700), lithium phenyl-2,4,6-trimethylbenzoylphosphinate (LAP) with purity ≥ 95%, were purchased from Sigma-Aldrich (St. Louis, MO, United States). Dialysis membrane (Mw cut-off (MWCO): 14 kDa) was obtained from Membra-Cel™, United States. Dyes: Congo Red and Methylene Blue were purchased from TCI (TCI EUROPE N.V., Zwijndrecht, Belgium).

Methacrylated gelatin was synthesized following our previous protocols [22, 34].

### 4.2 Resin formulation and volumetric printing

Volumetric printing was employed to create bioresin-based disks and constructs with various geometrical designs, including straight channels and angular configurations (60°, 90°, and 110°). Constructs were fabricated using a Gel-PEG resin, optimized for mechanical stability and biocompatibility (data not shown). The resin was based on GelMA 5 % (w/v), PEGDA (10 % (v/v), and LAP 0.3 mg/mL, abbreviated as Gel-PEG. The control resin used was GelMA 10% (w/v). The resins were sterilized using a 0.22 µm Corning® bottle-top vacuum filter system prior to printing. The resins were dispensed into cylindrical borosilicate (BK7) glass vials (ø 15 mm) [35]. Vials were loaded into a commercial volumetric 3D printer (Tomolite, Readily3D, Switzerland; multiwavelength). The samples were thermally gelated at 4°C prior to printing. Straight channels and angular designs were designed (SolidWorks, St. Waltham, United States) and generated .STL files were loaded into the printer software (Apparite, Readily3D, Switzerland). Resin’s refractive index was measured (SmartRef, LAB Meister, Anton Paar GmbH, Graz, Austria) and inputted into the software. Printing was performed with a 405 nm light source. After printing, the vials were heated to 37°C, uncured resin was collected and printed constructs were washed gently with 37 °C PBS [11, 36]. Printed constructs were rinsed with sterile phosphate-buffered saline (PBS) to remove residual resin and stored in PBS until further use. The printed constructs were characterized using brightfield microscopy to evaluate structural integrity, geometric fidelity and perfusion capabilities. Perfusion tests were conducted by injecting red food dye into the channels, assessing flow uniformity and potential obstructions.

### 4.3 Characterization

#### Rheological properties

Photocrosslinking of the resins was assessed Using an Anton Paar, MCR 302 rheometer (Anton Paar, Ghent, Belgium). Time sweep experiments were performed at a frequency of 1.0 Hz, with 1.0% constant strain at 25°C (n = 3, independent measurements); 15 seconds after the start of the measurement the light source (Dymax, VisiCure – 405 nm, Mavom, Kontich, Belgium) was activated for the remaining 105 s. The plate-plate measuring system was used, with a 25 mm diameter upper plate and a gap size of 100 µm [10, 37, 38].

#### Microscopic evaluations and Holographic imaging

Brightfield images during cell culture were taken using a Cell Imager (EVOS XL Core Configured Cell Imager, Invitrogen, Waltham, Massachusetts, United States). Fluorescently labelled samples were observed under a Leica Thunder DMi8 Inverted LED Fluorescence Motorized Microscope (Wetzlar, Germany). Micrographs were randomly taken during the observation [39].

Cell adhesion sites to the substrate material were observed using holotomography imaging on live cells (Tomocube, HT-X1, Daejeon, Republic of Korea). The reference refractive index was set at 1.337. Multiple z-stack images were taken and reconstructed to investigate cell attachment and morphology on different substrates.

To evaluate shape fidelity, we designed a multi-channel construct with three parallel channels of 500 µm, 700 µm, and 1000 µm diameters using SolidWorks. The model was exported as an .STL file and printed using three hydrogel formulations: Gel-PEG resin, commercial GelMA (60 % degree of substitution), and in-house synthesized GelMA (40 % degree of substitution). Printed constructs were stored in PBS at 37 °C and imaged on days 0, 1, 3, and 7 using a Zeiss AxioObserver microscope with Coherence Contrast (C-DIC) mode to visualize internal channel geometry. Channel diameters were measured at multiple points along their length, and average values were calculated. Dimensional changes were reported as a percentage relative to day 0 to assess structural stability over time.

Shape fidelity of disks was evaluated on volumetrically 3D printed hydrogels using an electronic hand caliper and compared to the theoretical value (digital input into the volumetric 3D printer).

#### Material swelling

The swelling properties of the hydrogels with and without B and TA treatment were investigated (n=3) [40]. The hydrogels were first freeze-dried, weighted, and then immersed in 3 mL of PBS (1X) in a sealed 24-well plate, and kept at 37°C. After 15 min, 30 min, 1h, 2h, 4h, and 6h of incubation in PBS, the samples were removed and excess PBS was drained with a paper towel, then the sample’s mass was recorded. The swelling ratio was calculated according to Equation 1 (**Eq. 1**).

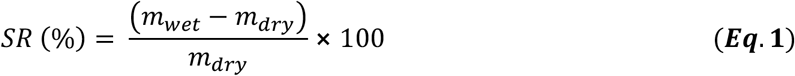

#### Mechanical properties

The evaluation of the stiffness of GelMA and Gel-PEG hydrogels was carried using a Chiaro Nanoindenter (Optics11Life). Resins were prepared and hydrogels were 3D printed as described above. Disks of GelMA and Gel-PEG were immersed in PBS and probed (Canteliver Geo factor in air: 3.42; stiffness: 0.014 N/m; Tip radius: 26.5 µm) with a load of 0.08 µN at 30 µm/s. For all the samples, the Young’s moduli were obtained using Hertzian contact curve fit model within the 4-7 µm indentation range.

Hydrogel viscoelastic properties were evaluated using the ElastoSens™ Bio2 system (Rheolution, Inc., Montreal, QC, Canada) at 37 °C. For polymerization kinetics, 2.0 mL of uncrosslinked hydrogel resin was added to sample cups immediately after degassing. Measurements were conducted in stiff test mode at 50% light intensity, with time-resolved storage modulus (G′) recorded *in situ* during photo-crosslinking. For long-term stability assessment, 2.0 mL of pre-crosslinked hydrogel was loaded into detachable sample holders and mounted on the instrument. Free resonance measurements were performed within the linear viscoelastic regime for 5 min. Between time points, samples were incubated in PBS at 37 °C. PBS was removed prior to each measurement and G′ values were recorded over seven days to monitor mechanical stability [41, 42].

#### Cell culture and seeding

Human osteosarcoma cell line 143b was maintained with Dulbecco’s minimal essential medium (high glucose), supplemented with 10 % FBS, 100 µg/mL Penicillin-Streptomycin (Thermo-Fisher Scientific, Massachusetts, United States), in a humidified CO_2_ chamber (37°C, 5 % CO_2_).

Human Umbilical Vein Endothelial Cells (HUVECs) purchased from Lonza (Basel, Switzerland) were maintained with Human Large Vessel Endothelial Cell Basal Medium (high glucose), supplemented with, 1 % Penicillin-Streptomycin and 0.1 % Low Serum Growth Supplement (LSGS) (Thermo-Fisher Scientific, Massachusetts, United States), in a humidified CO_2_ chamber (37°C, 5 % CO_2_).

To evaluate hydrogel biocompatibility and ability to serve as a support for cell growth and proliferation, cells were cultured on the hydrogel discs (seeding density 50.000/cm^2^) and daily monitored.

#### Channel seeding

After preincubation of 8-24 h in the appropriate cell medium, vascular models were removed from the incubator. The medium was aspirated and a 200-μL pipette was used to gently wash out the medium out of the channels, filling them with air instead [43]. Cells were dispersed at a seeding density of 2 million/mL and the channels were gently filled with 20-25-μL of the cell suspension ensuring that the air is fully pushed out of the channels. Once the seeding was complete, the vascular models were returned to the incubator. After 24-30 h of incubation, non-adherent cells and cell debris were removed by gently perfusing the channels with fresh medium. Samples were cultured for one week with the media refreshed every day.

#### Cell staining

To evaluate cellular morphology and distribution within volumetrically printed channels cells were fixed with 4% paraformaldehyde for 30 min at room temperature, and permeabilized with 0.3% Triton X-100 (Sigma-Aldrich, 93443) 0.5% Tween and 0.5% sodium azide in 1% Bovine Serum Albumin (BSA, Sigma-Aldrich, A2153) in phosphate buffered saline (PBS, Sigma-Aldrich, P4417) at room temperature for 5 min. Next samples were washed with 1% BSA/PBS and kept at 4°C or directly used for staining.

Cytoskeleton staining (F-actin) was performed with ReadyProbes™ Reagent F-Actin Phalloidin Conjugates (Thermo-Fisher Scientific, Massachusetts, United States). Following the manufacturer’s instructions 2 droplets were used per 1 mL of staining solution and 5 μg/ml DAPI (Sigma-Aldrich, D9542) was added to stain nuclei.

For immunostaining primary Rabbit antibody anti-CD31, 1:500 (Novus Biologicals, NB100-2284) for HUVECs and Rabbit anti-H3K27me3 1:500 (Thermo Fisher Scientific, MA5-11198) for 143b cells were diluted in 0.1% Tween-20, 0.05% sodium azide in 1% BSA/PBS and incubated in a humid chamber (4^°^C, overnight). Triplicate washes of 0.1% Triton-X in 1% BSA/PBS were carried out (room temperature, 10 min). Secondary antibody Donkey anti-Rabbit-555 1:1000 (Thermo Fisher Scientific, A32816) was diluted in 0.1% Tween-20, 0.05% sodium azide in 1% BSA/PBS, with 5 μg/ml DAPI (Sigma-Aldrich, D9542) and incubated in a humid chamber (dark, room temperature, 1 h). Triplicate washes of 0.1% Triton-X in 1% BSA/PBS were carried out (room temperature, 10 min) [20]. Immunostained samples were imaged with an inverted Leica Thunder DMi8 LED Fluorescence Microscope (Wetzlar, Germany). To evaluate cell density, we counted nuclei and the area they occupy in specified design regions. Images were post-processed and analyzed through ImageJ/FIJI [39].

### 4.4 Statistical analysis

All data are reported as mean ± standard deviation (SD), with the number of independent samples (n) indicated in the figure captions. All statistical analyses were performed using Origin Pro 2021 and Prism 10 (GraphPad).

Given the small sample sizes, non-parametric statistical methods were used to ensure the robustness of the analysis without reliance on normality assumptions. The Kruskal-Wallis test was employed for comparisons between groups in the analysis of cell number and occupied area in different geometrical designs, followed by Dunn-Sidak’s test for post-hoc pairwise comparisons.

## Author Contributions

Conceptualization (JS, PT, AS, SJH), Data curation (JS, PT), Formal Analysis (JS, PT), Investigation (JS, PT), Methodology (JS, PT, AS, SJH), Project administration (AS, SJH), Resources (JS, AS, SJH), Supervision (PT, AS, SJH), Validation (JS, PT), Visualization (JS, PT), Writing – original draft (JS, PT), Writing – review & editing (JS, PT, AS, SJH).

All authors have read and agreed to the published version of the manuscript.

## Institutional Review Board Statement

Not applicable.

## Informed Consent Statement

Not applicable.

## Acknowledgements

The graphical abstract and Figures schematics were created using BioRender.com.

J.S.S. gratefully acknowledges the support of a grant from the FNRS (J.S.S. FNRS-Aspirant, Grant No. FC 46599).

This work was supported by the ULB–UNIL privileged partnership program.

A remotely controlled perfusion chamber with a pressure sensor and two peristaltic pump units was designed and assembled by A4BEE Sp. z o.o. (Wrocław, Poland). We thank the team of engineers from A4BEE Sp. z o.o. for their technical support in the preparation of the microfluidic modules and in particular we thank Kinga Surmacz and Paweł Godawa for their valuable discussions.

## Data Availability Statement

The authors declare that part of the data supporting the findings of this study is provided as supplementary information. Additional data may be available from the corresponding author upon request.

