## Supplemental files for "Geometrical Designs in Volumetric Bioprinting to Study Cellular Behaviors in Engineered Constructs"

SUPPLEMENTARY INFORMATION

***S1 Results***

***S1.1 Indentation testing***

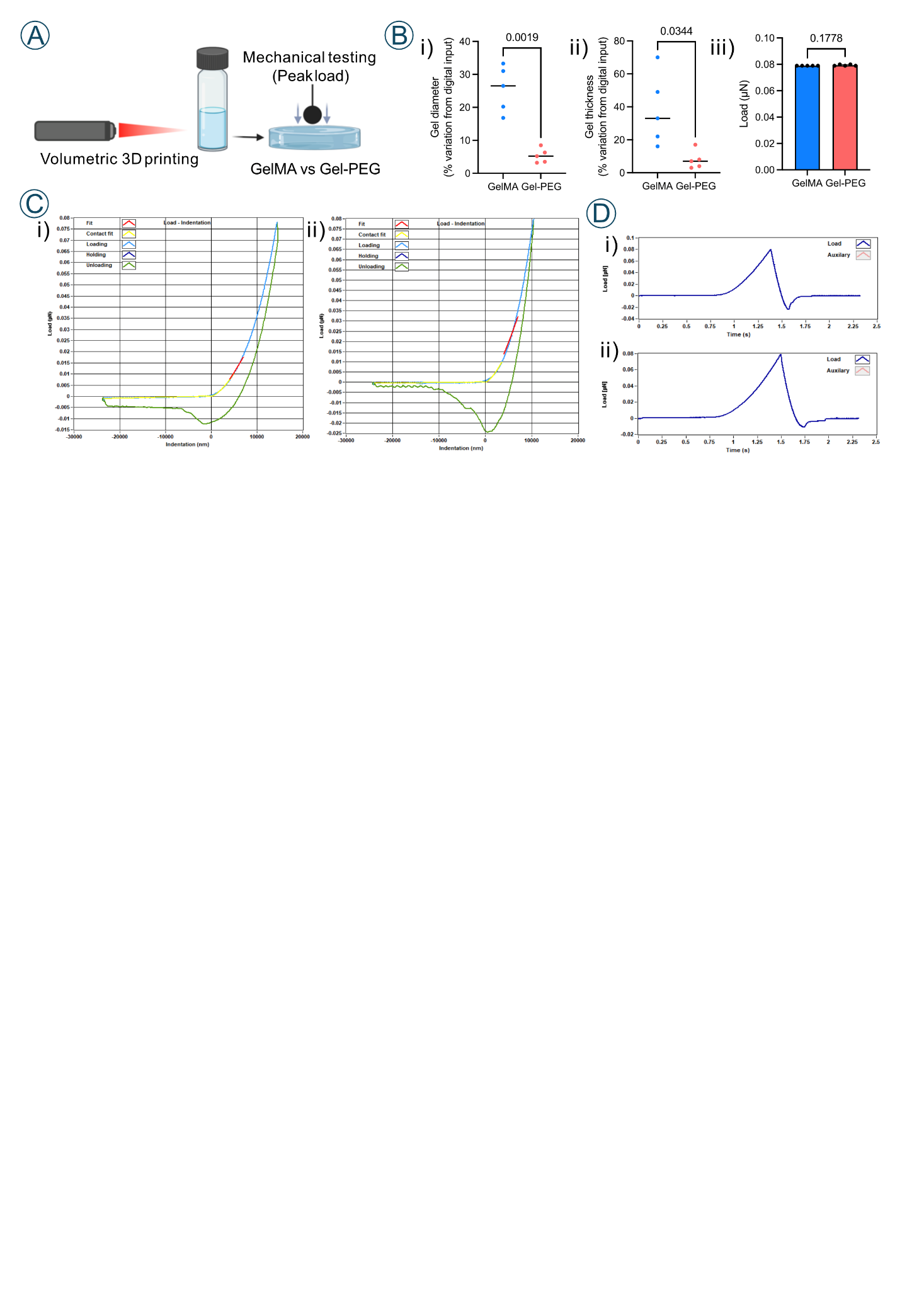

***Figure S1*** *Comparison of printing fidelity and mechanical properties of GelMA and Gel-PEG hydrogels. A) Schematic of experimental workflow: volumetric 3D printing of GelMA and Gel-PEG hydrogels followed by mechanical testing (peak indentation load); B) Quantification of 3D printing fidelity showing percent deviation from digital input for gel diameter (i) and thickness (ii). Gel-PEG hydrogels exhibited significantly smaller deviations than GelMA, indicating better dimensional accuracy. N = 5 gels per group. Statistical test: unpaired two-tailed t-test; Applied peak loads for Gel-PEG and GelMA hydrogels were similar between groups (iii) N = 5 gels per group, n ≥ 16 measurements per gel. Statistical test: Welch’s t-test. C) Representative load–indentation curves for GelMA (i) and Gel-PEG (ii) hydrogels, with phases of loading (blue), and unloading (green). The Hertzian contact curve fit model was used for Young’s modulus calculation within 4-7um indentation depth (red). D) Representative load–time profiles for GelMA (i) and Gel-PEG (ii) during indentation tests.*

***S1.2 Holographic microscopy images showing cell attachment***

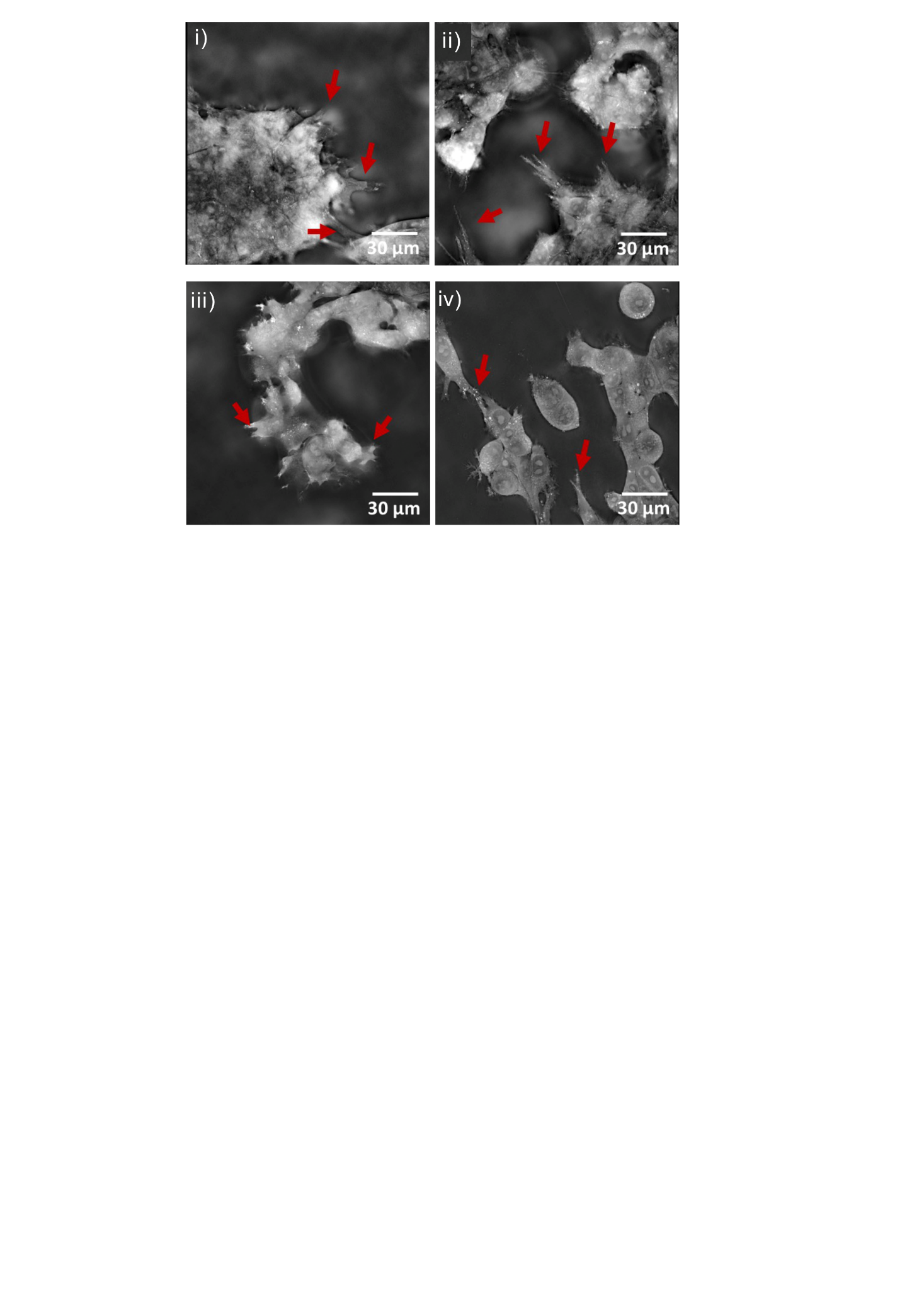

***Figure S2*** *Holographic microscopy images showing cell attachment to the surface of different materials (i) 143b cells on GelMA surface, (ii) 143b cells on GelMA-PEGDA surface, (iii) HUVECs on GelMA surface, (iv) HUVECs on GelMA-PEGDA surface. Red arrows point to the cell adhesion points.*

***S1.3 143b cells lining the volumetrically printed channels***

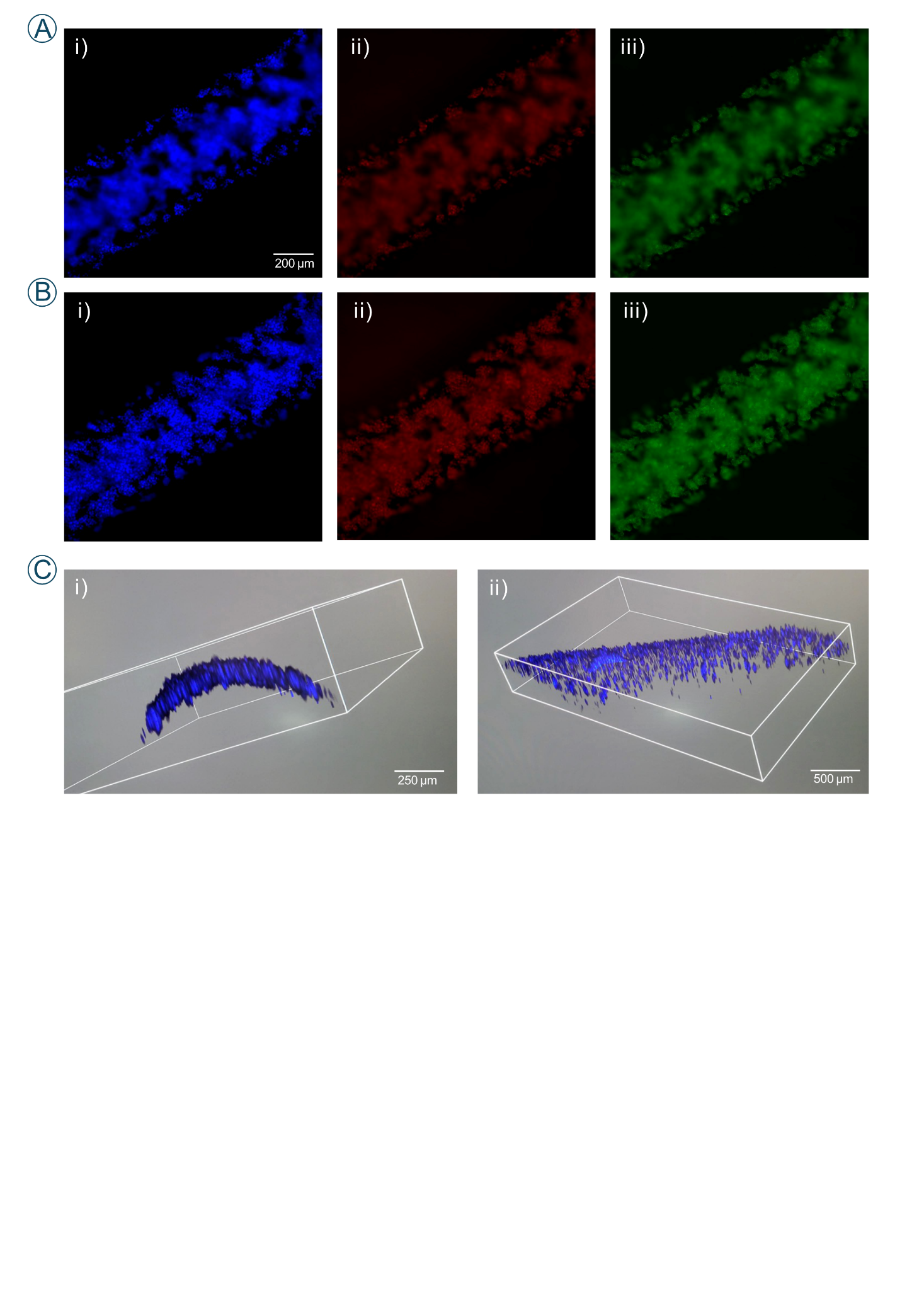

***Figure S3*** *Fluorescently labelled 143b cells cultured within geometrical hydrogel confinements for 7 days* ***A)*** *focus on the channel edges, B) focus on the bottom of the channel, blue corresponds to cell nuclei (i), red to expressed histone marker (H3K27me^3^) (ii) and green to cytoskeleton (iii); C) DAPI-blue labelled cell nuclei patterning the channel geometry side view (i), top view (ii).*
